# Sexual conflict drives micro- and macroevolution of sexual dimorphism in immunity

**DOI:** 10.1101/2020.12.20.423717

**Authors:** Basabi Bagchi, Quentin Corbel, Imroze Khan, Ellen Payne, Devshuvam Banerji, Johanna Liljestrand-Rönn, Ivain Martinossi-Allibert, Julian Baur, Ahmed Sayadi, Elina Immonen, Göran Arnqvist, Irene Söderhäll, David Berger

**Affiliations:** Department of Biological Sciences, Ashoka University; Department of Ecology and Genetics, Program of Animal Ecology. Evolutionary Biology Centre, Uppsala University; Cavanilles Institute of Biodiversity and Evolutionary Biology, University of Valencia; Department of Ecology and Genetics, Program of Evolutionary Biology. Evolutionary Biology Centre, Uppsala University; Department of Organismal Biology, Evolutionary Biology Centre, Uppsala University; Department of Chemistry, Biochemistry, Uppsala University; Department of Organismal Biology, Program of Comparative Physiology. Uppsala University

**Author notes:** Equal contribution and presented in alphabetical order (Basabi Bagchi, Quentin Corbel, Imroze Khan, Ellen Payne).

**Keywords:** immunity, sexual selection, sexual conflict, trade-off, mating, sexual dimorphism, sexually transmitted disease, phenoloxidase, experimental evolution, *Callosobruchus maculatus*.

## Abstract

**Background:** Sexual selection can have major effects on mating rates and sex-specific costs of mating and may thereby influence sex-differences in immunity as well as associated host-pathogen dynamics. Yet, experimental evidence linking the mating system to evolved sexual dimorphism in immunity are scarce and the direct effects of mating rate on immunity are not well established. Here, we use transcriptomic analyses, experimental evolution and phylogenetic comparative methods to study the association between the mating system and sexual dimorphism in immunity in seed beetles, where mating causes internal injuries in females.

**Results:** We demonstrate that female phenoloxidase (PO) activity, involved in wound healing and defence against parasitic infections, is elevated relative to males. This difference is accompanied by concomitant sex-differences in the expression of genes in the pro-phenoloxidase activating cascade. We document substantial phenotypic plasticity in female PO activity in response to mating and show that experimental evolution under enforced monogamy (resulting in low remating rates and sexual conflict relative to natural polygamy) rapidly decreases female (but not male) PO activity. Moreover, monogamous females have evolved increased tolerance to bacterial infection unrelated to mating, implying that female responses to costly mating may trade off with other aspects of immune defence, an hypothesis which broadly accords with the documented sex differences in gene expression. Finally, female (but not male) PO activity shows correlated evolution with the perceived harmfulness of male genitalia across 12 species of seed beetles, suggesting that sexual conflict has a significant influence on sexual dimorphisms in immunity in this group of insects.

**Conclusions:** Our study provides insights into the links between sexual conflict and sexual dimorphism in immunity at the molecular and phenotypic level and suggests that selection pressures moulded by mating interactions can lead to a sex-specific mosaic of immune responses with important implications for host-pathogen dynamics in sexually reproducing organisms.

## Introduction

Sex differences in immunity are widespread across animal taxa (1–4) and are believed to reflect sex-specific selection and sexually dimorphic life histories (5–13). Sexual dimorphism in immunity may have important consequences both for sex-specific rates of reproduction and survival, with potential impact on population demography (14–18), and for the spread of pathogens. For example, distinct male and female immune systems present more diverse host targets (1,19,20) and this may influence both disease transmission, infection rates and the expression and evolution of pathogen virulence (5,16–18,21–29).

Investment in immune defence is costly. These costs have most often been observed as reductions in fecundity, effectively translating into reproduction-survival trade-offs in the presence of pathogens (9,10,12,22,30–34). In polygamous species, where sexual selection on males is intense, females are often predicted to gain more than males from investing in survival and longevity at the cost of current reproduction and mating effort (3,9,35) and are therefore also predicted to invest more in immunity than males (but see: (2,10,24,35–37)). Sexual selection may also have pronounced direct effects on optimal investment in immunity, as it may dictate the economics of reproduction (23,27,38,39) and lead to elevated mating rates (40), which in turn may increase disease transmission (16,24,25,28). Indeed, it has been suggested that sexual dimorphism in immunity should increase with sex-differences in optimal mating rates and the strength of sexual selection (5,13,21,23,27,41,42).

The effects of sexual selection on sex-differences in immune investment may be magnified in systems where mating is harmful for females, through costs such as the transfer of pathogens during mating, transfer of immunosuppressive seminal fluid substances, or direct physical injury (23,28,43–45). Such male-imposed mating costs are believed to be results of sexual conflict driven by the different evolutionary interests of the sexes (6–8,46), in which male adaptations evolve to increase reproductive success in competition with other males despite impairing the health of their female mating partners. Females, in turn, evolve counter-adaptations to alleviate the harm inflicted by males resulting in a coevolutionary arms race between the sexes (23,43,46,47). Female immune responses may represent one type of such counter-adaptation (23,27,48,49). This suggests that infections or harm on females, induced by sexually selected male mating strategies, may be a significant selection pressure on female immunity in polyandrous taxa (21,24,27,41,50). Hence, the evolution of sexual dimorphism in immunity may in part be a result of male-imposed costs of mating in females.

Yet, whether and how sexual conflict, or just mating *per se,* affect tissue-specific and general immunity in the sexes is not well understood (5,22,23,44,51). It has, for example, been suggested that tissue-specific (i.e. in the reproductive tract) immune responses upon mating can lead to allocation trade-offs with systemic immunity (44, 52), but few studies have provided direct experimental evidence for a causal link between the mating system and the evolution of sex-specific immunity trade-offs (2,37,49,50,53). To fill this empirical void, we assessed how variation in the intensity of sexual conflict and mating rates in the seed beetle *Callosobruchus maculatus* affects i) the evolution of male and female phenoloxidase (PO) activity, a major component of invertebrate immunity involved in wound healing and encapsulation of pathogens (54, 55), and ii) associated immunopathological consequences of bacterial infections unrelated to mating.

Sexual selection is intense in *C. maculatus*, including both pre- and post-copulatory processes (56–61), leading to sexual conflict over optimal mating rate and to male traits that cause harm in females during mating (59,60,62,63). The male genitalia carry spines and males with longer spines have greater fertilization success but the spines cause internal injuries in females during mating, leaving females with melanized scars in the reproductive tract as a result of the wound-healing process (59,60,62). Injurious copulations are wide-spread in insects and may serve several functions, with the ultimate aim to increase male competitive fertilization success (64, 65). This may select for increased immune defence locally in the female reproductive tract to enable efficient wound healing and limit female susceptibility to sexually transmitted pathogens (66). Here, we show that PO activity in *C. maculatus* females is high (see also: (48)) and responds dynamically to mating, while it is very low in males. These sex differences are also mirrored in the expression of several key genes regulating PO activity and related immune reactions. Experimental removal of sexual selection and conflict led to rapid laboratory evolution of decreased female (but not male) investment in PO activity. These changes were accompanied by the evolution of increased female tolerance to bacterial infection unrelated to mating, suggesting a trade-off between female responses to harmful mating and tolerance to other infections. The PO response was paralleled at a macroevolutionary scale, signified by correlated evolution between male genital morphology and sexual dimorphism in PO activity across 12 species of seed beetles.

## Results

### Mating status and sex-biased gene expression in the prophenoloxidase-activating cascade

The prophenoloxidase (proPO) activating cascade leads to the production of active PO, which serves as an important defence in invertebrates against pathogenic bacteria, fungi and viruses (54,55,67,68). Additionally, proPO has been implicated in cuticle tanning and other developmental processes, as well as reproduction (reviewed in: (54,55,68)). PO aids in wound healing and encapsulation of parasitic infections, and killing of pathogens by generation of toxic secondary metabolites, such as reactive oxygen species (54,55,68–73). However, the production of PO is strictly regulated (74, 75) as it is both energetically costly and the generation of toxic secondary metabolites can cause self-harm via immunopathological responses (68,73,76,77), predicting that investment in PO activity could incur costs to other fitness related traits (22,55,68,73). In Figure 1a we delineate the general hypothesis for relationships between key components of the proPO cascade based on functional annotations in insects and other invertebrates (reviewed in: (54,55,67,68,78)). To gain insights into how sexual selection and conflict may affect investment in PO and other correlated immunity traits in *C. maculatus*, we explored sex-biased gene expression of five orthologs mapping to sequences of proteins functionally annotated for these key components (Supplementary Table 1, Figure 1b). Spätzle processing enzyme (SPE) is involved in the processes that cleaves proPO into active PO. We found that the expression of the *C. maculatus* orthologs of both SPE and proPO are significantly female-biased in virgin adults. Mating increased transcription of proPO in males leading to similar expression levels in the sexes, whereas expression of SPE tends to increase in both sexes post mating and remains female-biased (Figure 1b, Supplementary Table 1). These results suggest that females invest heavily in PO activity via SPE and proPO. SPE also initiates the modification of spätzle (SPZ) and downstream TOLL-regulated antimicrobial peptides (AMPs), which offer inducible immunity to pathogens. This may thus set the stage for a trade-off between PO (encapsulation and wound healing) and SPZ (AMP-production) (see: e.g. (79, 80) (Figure 1a). Overactivation of the proPO cascade may also lead to the production of toxic secondary metabolites (68, 73), suggesting that excessive signalling via SPE to produce high levels of both SPZ and PO may come at a cost to overall health (76, 77). Interestingly, production of serine protease inhibitors (serpins) via the TOLL-pathway exerts negative feedback and control over the proPO cascade (81), and orthologs of both SPZ and the two putative serpins that we identified in *C. maculatus* had strong male-biased expression (Figure 1b, Supplementary Table 1). These patterns in gene expression thus suggest a putative functional basis for sex-specific immunity via the pro-PO activating cascade, where we hypothesize that females (relative to males) should invest more in PO activity in their reproductive tract in response to harmful mating and the need for wound healing, but that this investment might come at the potential cost of reduced AMP-production and/or toxic side-effects of overactivation of the proPO cascade.

**Figure 1:**
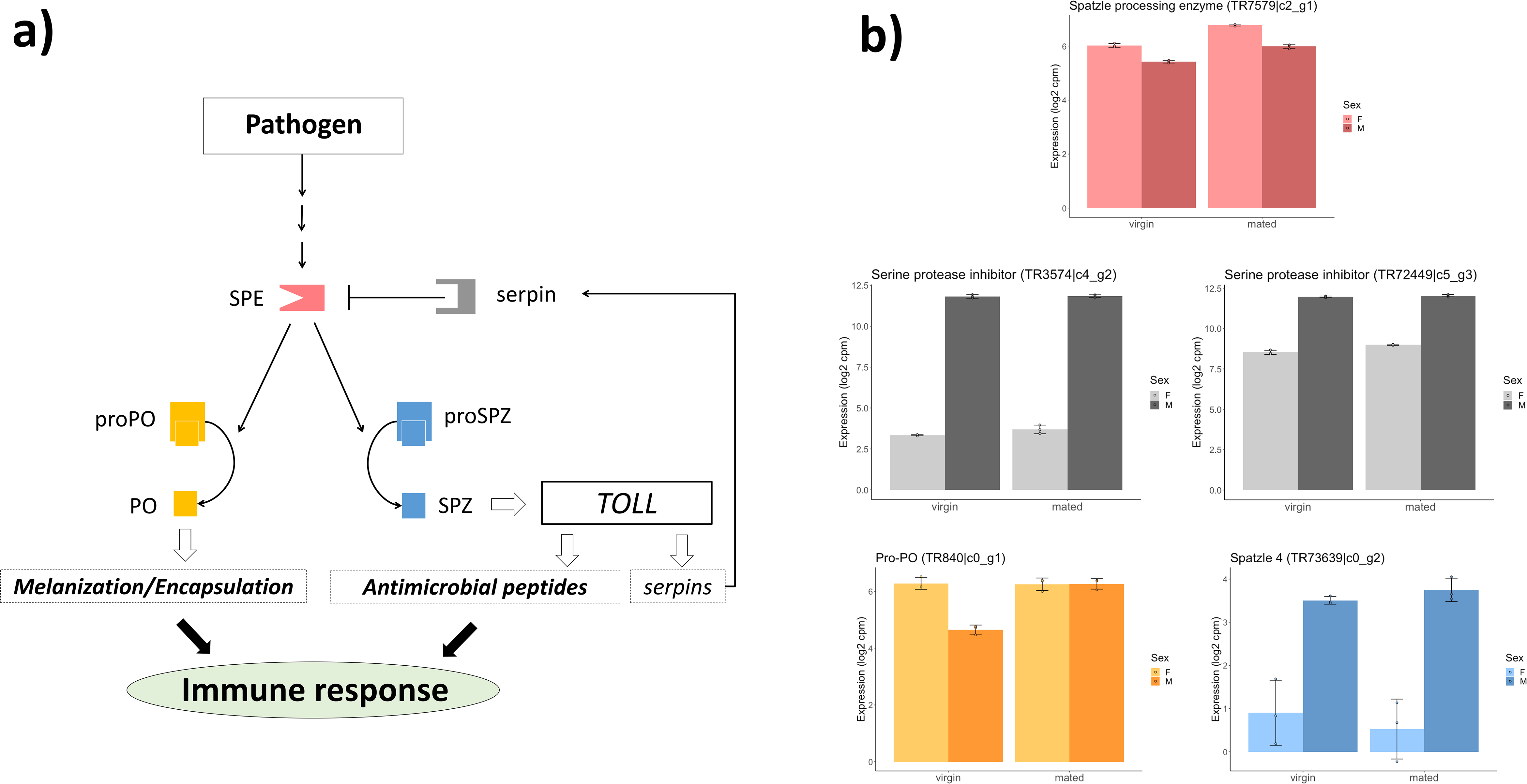
Sex-biased gene expression in the proPO signalling cascade. In **a)** schematic representation of key proteins in the proPO activating cascade, based on previous studies of insects and other invertebrates (reviewed in: (54,55,67,73,78)). In **b)** sex-bias and effects of mating status on gene expression in the abdomen for *C. maculatus* orthologs from published data (128) mapped to the sequences of the functionally annotated proteins. Full results in Supplementary Table 1. Spätzle processing enzyme (SPE: pink) initiates cleavage of proPO (yellow) into active PO, which ultimately leads to wound healing as well as encapsulation and killing of foreign pathogens. However, SPE also regulates the production of Spätzle protein (SPZ) from proSPZ (blue), which ultimately leads to increased production of antimicrobial peptides (AMPs) via the TOLL pathway, which offers inducible immunity against pathogens, thus setting the stage for an allocation trade-off between PO-activity and AMP-production. Overactivation of the proPO cascade has toxic side-effects via the production of secondary metabolites, suggesting that overproduction of SPE may come at a cost to overall health. Here, production of serine protease inhibitors (serpins: grey) in the TOLL-pathway exerts negative feedback and control over the cascade. **b)** *C. maculatus* females show higher expression of SPE (pink) and proPO (yellow) as virgins. Males show higher expression of proSPZ (blue) and serpins (grey). These patterns in gene expression suggest a mechanistic basis for sex-specific immunity trade-offs between different components in the pro-PO activating cascade, where females are predicted to invest more in PO-activity (wound healing and potentially encapsulation of pathogens transferred at mating) in their reproductive tract in response to mating, at the potential cost of reduced inducible immunity via AMP-production and/or toxic side-effects of overactivation of the proPO cascade.

### Sex-specific regulation of phenoloxidase activity

We measured PO activity in homogenized whole-body samples of male and female larvae, pupa and adults. The three life stages showed significant differences in mass-corrected PO activity averaged across the sexes (F_2,33_ = 17.7, p < 0.001, Figure 2a). Some larvae showed detectable levels of PO activity. Since we could not determine the sex of the larvae, sex-differences in the larval stage can neither be confirmed nor rejected. Neither male nor female pupae showed measurable levels of PO activity, whereupon there was a drastic and female-limited up-regulation in the virgin adults. Strikingly, virgin males did not show any PO-activity, which was also the case for mated males (see further below), despite clear expression of the proPO gene in males, especially following mating (Figure 1b). It seems that proPO is not converted to PO in males to the same extent that it is in females, and other proteins such as proPO activating factors (PPAFs), for which we could not confidently identify gene transcripts, might be involved in regulating sex differences in how proPO is converted into active PO. The observed effect size of sex on PO activity in virgin adults was, Hedges’ *g* = 2.08, which is high relative to what is typical in insects (mean Hedges’ *g* = 0.55; see (2)) and for animals in general (mean Hedges’ *g* = 0.39; see (2)).

**Figure 2:**
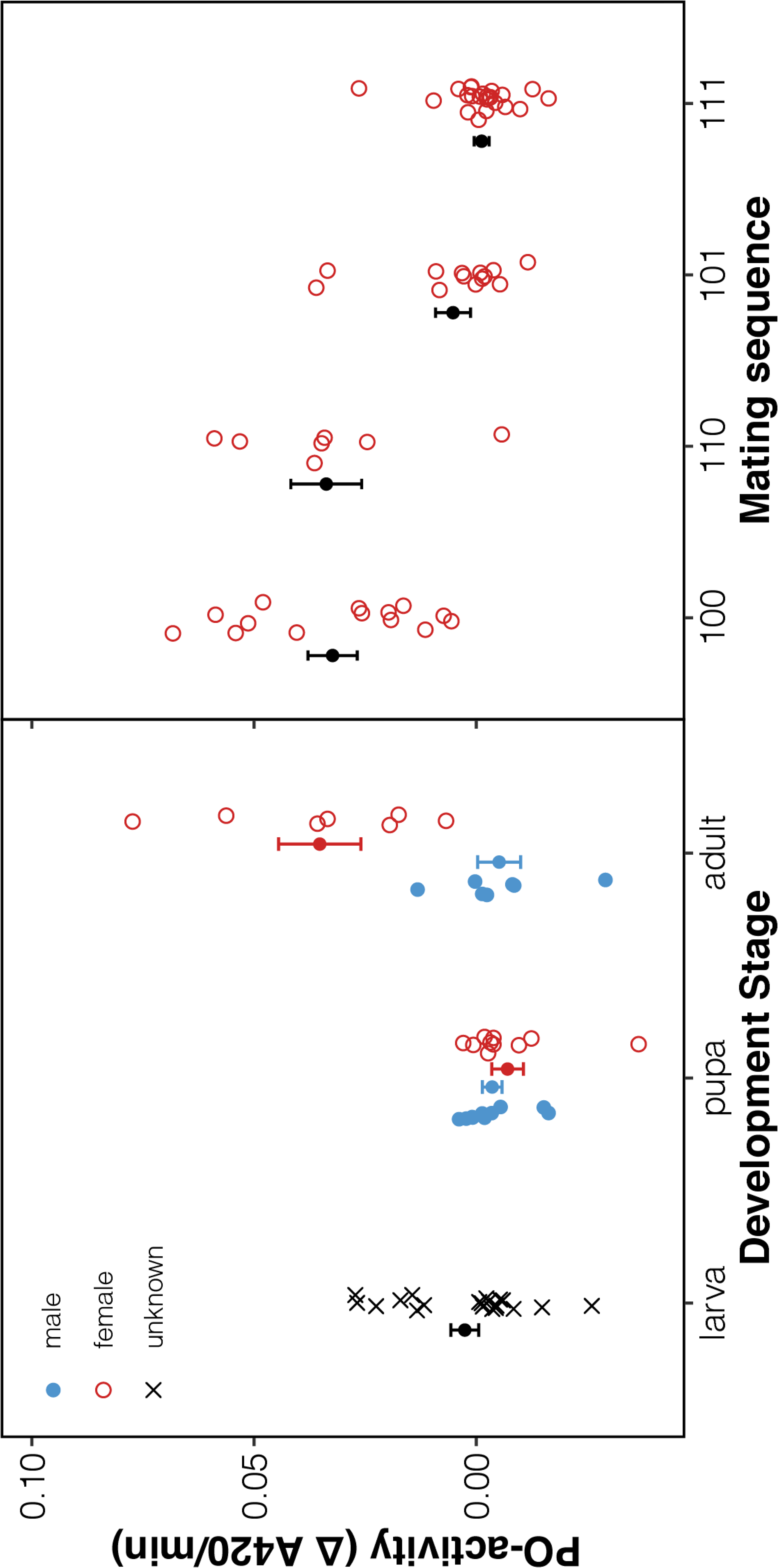
Sex-specific regulation of phenoloxidase levels. **(a)** There were significant differences in PO activity throughout development, with levels near zero detected in male (blue) and female (red) pupae and virgin adult males, but detectable levels in (unsexed = black) larvae and high levels in virgin adult females. **(b)** PO activity measured on day 3 in females mated only on day one (100), day one and two (110), day one and three (101), or on all days (111) (open symbols). A second experiment measured PO activity for a random set of females assigned to treatments 100 and 001 (mated only on day three) (filled symbols). Female PO activity is reduced after mating but is then quickly restored (compare also to virgin females (i.e. 000 treatment) in **(a)**. Shown are means ± 1 SE and individual observations. PO-activity was corrected for body mass by including mass as a covariate in all analyses but is here displayed as raw data since the mean amount of tissue in samples was similar for all groups.

To further understand the function of the female-bias in adults, we explored how female PO activity responds to mating. We mated females either only on day one of adult life (treatment 100), on day one and two (110), on day one and three (101), or on all days (111) and measured levels of PO activity subsequently on the third day (2h post mating in 101- and 111 females; ca. 24 and 48h post mating in 110- and 100 females, respectively). The differences among the four treatments were substantial (F_3,52_= 18.7, p < 0.001, Supplementary Table 2). The PO activity was high in females when some time had elapsed between mating and PO measurement (i.e. 100- and 110-females), while the levels were near zero when PO activity was measured directly after mating (i.e. 101- and 111 females) (Figure 2b). The treatment groups described above represent non-random samples of females, as not all females can be made to remate on a given day. We therefore also conducted a second experiment with random samples of 100 and 001 females. This showed that the two treatments differed significantly (Mann-Whitney U test: W = 15, p < 0.001); 001 females had PO activity close to zero similar to 101 and 111 females in the first experiment, whereas 100 females had high PO activity similar to virgins, 100 and 110 females of the first experiment (Figure 2). Hence, female PO activity decreases after mating but can be rapidly recovered to initial levels post mating. These results accord with the observed female upregulation of SPE in response to mating and the unconditionally high expression of proPO in the female abdomen (Figure 1b, SI Table 1).

Using a subset of 25 females from the same population and same generation as the first experiment, we performed a subsequent analysis of PO activity in oviposited eggs. This analysis showed that decreases in female PO activity following mating is not due to PO investment in offspring, as all five samples of pooled eggs showed very low (undetectable) levels of PO activity, despite each sample representing about half of the lifetime egg production of a single female. We found no evidence of a reproduction-immunity trade-off as there was no relationship between the number of eggs laid by the females over the two days of the first experiment and their subsequent measure of PO activity (Supplementary Table 2). Although immunity-reproduction trade-offs are readily observed in insects (9,12,13,22), PO investment does not always correlate negatively with fecundity (e.g. (82, 83)). Moreover, variation in overall phenotypic and genetic condition (84, 85), as well as the amount of male harm inflicted on females (86), could have masked a putative trade-off. Alternatively, trade-offs with PO investment could materialize for other life-history traits (9,30,87), and/or other components of immunity (22, 55) (see: Figure 1a and further below).

### Experimental evolution of phenoloxidase activity under different mating systems

To directly test the hypothesis that sexual selection and conflict over mating is causing the observed sexual dimorphism in immunity in *C. maculatus*, we compared the levels of PO activity in males and females from replicate experimental evolution lines maintained for 27 generations under one of three alternative mating regimes; natural *polygamy* (natural selection and sexual selection – multiple mating); enforced *monogamy* (natural selection but excluding sexual selection – single mating); and *male-limited selection* (applying sexual selection but relaxing natural selection–multiple mating but female coevolution to reduce male harm prevented). The lines are further described in the *Methods* section and in (63,88,89). We predicted that females from polygamous lines that had evolved under frequent multiple mating would invest more in PO activity than females from monogamous lines, while the male-limited lines reveal the extent to which female PO activity may change in the polygamous mating system via genetic correlation when selection acts mainly via males. We also tested whether the direct effect of mating and reproduction on PO activity had evolved under the different mating systems by for all lines comparing PO activity of virgin and socially naïve individuals to that of beetles allowed to mate and reproduce for 48 hours in groups of 5 males and 5 females prior to the PO measurements.

We analysed the effects of experimental evolution regime crossed by mating treatment in Bayesian mixed effect models using the MCMCglmm package (90) for R (91). Experimental evolution line replicates, crossed with mating treatment, were included as random effects (priors and model specification in Supplementary 3). The mating treatment decreased body mass relative to the virgin treatment, revealing a sizeable investment in reproduction by both sexes (SI Table 3a). While males did show an up-regulation of proPO gene expression in response to mating (Figure 1b), they did not have any detectable levels of PO activity (n = 354, SI Table 3c), confirming that PO investment is strongly female-biased in the adult stage in *C. maculatus* (63). In females (N = 358 assays), the mating treatment significantly decreased PO activity (ΔPO = −0.029 (−0.022; −0.037), P_MCMC_ < 0.001) but this effect was similar in the three selection regimes (all pairwise interactions P_MCMC_ > 0.6) (Figure 3). Importantly, evolution without sexual conflict under the monogamy regime had led to a general decrease in female PO activity relative to the polygamy regime (ΔPO = −0.010 (−0.002; −0.018), P_MCMC_ = 0.030), confirming a key prediction. The monogamy regime also showed lower levels of PO activity compared to the male-limited regime, where females had been kept under relaxed selection (ΔPO = −0.011 (−0.004; −0.020), P_MCMC_ = 0.012). Accordingly, the polygamy and male-limited regime had similar levels of PO activity (P_MCMC_ > 0.8, Figure 3). Thus, as the expected number of matings decreased to a single mating in the monogamy regime, the optimal female strategy was to decrease PO activity, in support of the hypothesis that PO investment is costly and likely trades off against other female fitness components (22,50,55,68,92). If immune defence is costly, a corollary from allocation theory is that polygamous females should invest in PO in relation to their total energy reserves and expected number of partners. In contrast, among monogamous females we expect the evolution of decreased condition dependence due to their reduced need for PO activity. This is also what we find; there was a positive relationship between female body mass and PO activity in polygamous lines (slope = 0.011 (0.005; 0.016), P_MCMC_ < 0.001), whereas this relationship was absent in monogamous lines (P_MCMC_ = 0.48), and this regime-difference in the condition dependence of PO investment was significant (Δslope = 0.007 (0.001; 0.013), P_MCMC_ = 0.026, Figure 3).

**Figure 3:**
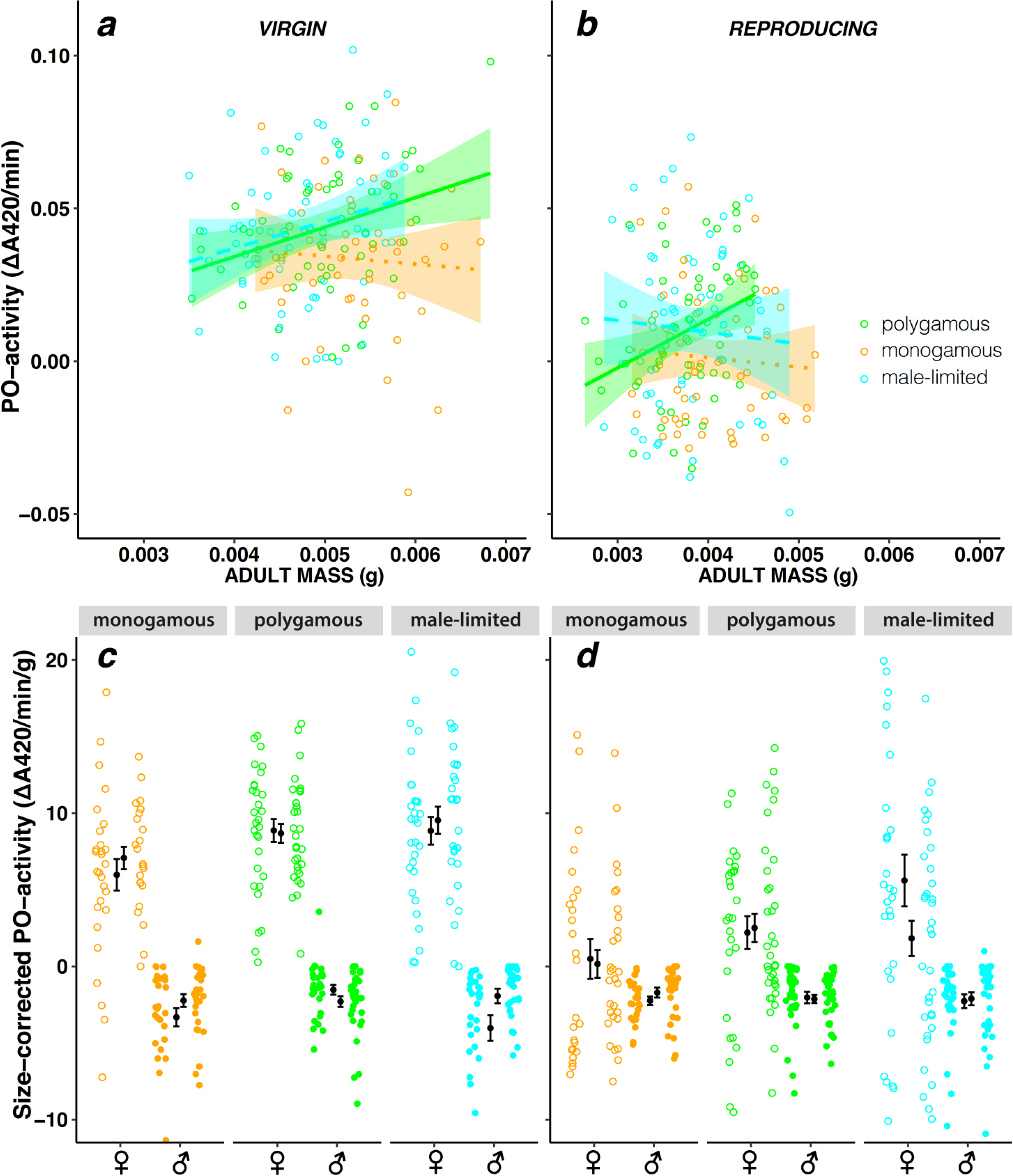
Microevolutionary change in PO activity during experimental evolution. PO activity measured from whole-body samples of virgin **(a)** and mated **(b)** females from polygamous (green) monogamous (orange) and male-limited (blue) evolution lines. The mating treatment significantly reduced female PO activity and male-limited and polygamous females had higher PO activity than monogamous females. Polygamous and monogamous females also differed significantly in the relationship between body mass and PO activity, suggesting that different allocation strategies evolved under the alternative mating regimes. Given are regression slopes, shaded 95% confidence limits, and individual observations. Males from the regimes did not express detectable levels of PO activity and showed no significant differences among regimes and mating treatments (Supplementary Table 1c). In the lower panels, sex differences in size-corrected PO activity is illustrated in each regime for **(c)** virgin and **(d)** reproducing beetles (means ± 1SE and individual measures).

Again, however, a fecundity cost of high PO activity was not apparent when comparing regimes; offspring production in the reproducing treatment was higher for females from the polygamy regime (showing higher levels of PO activity) than for monogamous females (with lower levels of PO activity) (SI Table 3b).

### Experimental evolution of the response to bacterial infection

To explore other possible immunological consequences of mating system and sexual conflict, which could be driven by trade-offs between investment in PO and other components of immunity (Figure 1), we measured survival in the monogamy and polygamy lines when exposed to bacterial infection in abdominal tissue adjacent to the reproductive tract. Females (total n = 1060, 24-48h past adult eclosion) were either virgin or mated prior to being infected with one of two doses (OD1 or OD2) of the entomopathogenic gram-positive bacteria, *Bacillus thuringiensis,* or a sham control (pricking with a sterilized needle dipped in PBS buffer). We analysed survival in mixed effects Cox proportional hazard models using the coxme package (93) for R, with regime and mating treatment as fixed effects and replicate lines as random effects. We also confirmed results by using the MCMCglmm package (90) to apply Bayesian mixed effect models on a binomial response variable (dead/alive on day 5 post infection), which allowed us to add fully crossed random effects (line by treatment) in the analysis (Full statistical summaries in Supplementary Tables 4a, b).

Females from the polygamy regime showed lower survival under bacterial infection compared to females from the monogamous regime (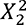 = 13.7, P = 0.001, Figure 4a-d). This result, albeit correlative, is in line with the hypothesis that the evolution of female immunity responses to expected harmful mating may trade-off against general susceptibility to infection. Mating by itself led to an increase in mortality (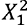 = 63.6, P < 0.001). However, there was no significant effect on susceptibility to infection of either mating status (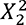 = 1.2, P = 0.56) or the interaction between evolution regime and mating status (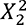 = 0.14, P = 0.93, Figure 4a-d). Although somewhat surprising, this result is not inconsistent with a trade-off between female PO investment in the reproductive tract and vulnerability to systemic infection caused by other pathogens, as also virgin females display high PO activity and high expression of genes in the proPO cascade prior to being mated (Figures 1b & 2). Virgin males from monogamous and polygamous regimes (which do not seem to invest in PO at all) did not show any strong differences in their response to bacterial infection (assessed in a separate experiment, 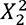 = 0.94, P = 0.63, Figure 4e,f). However, although we analyzed the same number of evolution lines in the male experiment, the total number of individuals analyzed was smaller (n = 270 for virgin males compared to n = 493 for virgin females), limiting direct comparisons between the male and female assays. Nevertheless, the male experiment did reveal an overall effect of the bacterial injection (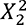 = 7.77, P = 0.021) and significantly greater survival of polygamous males (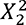 = 6.63, P = 0.010) (SI Table 4c).

**Figure 4:**
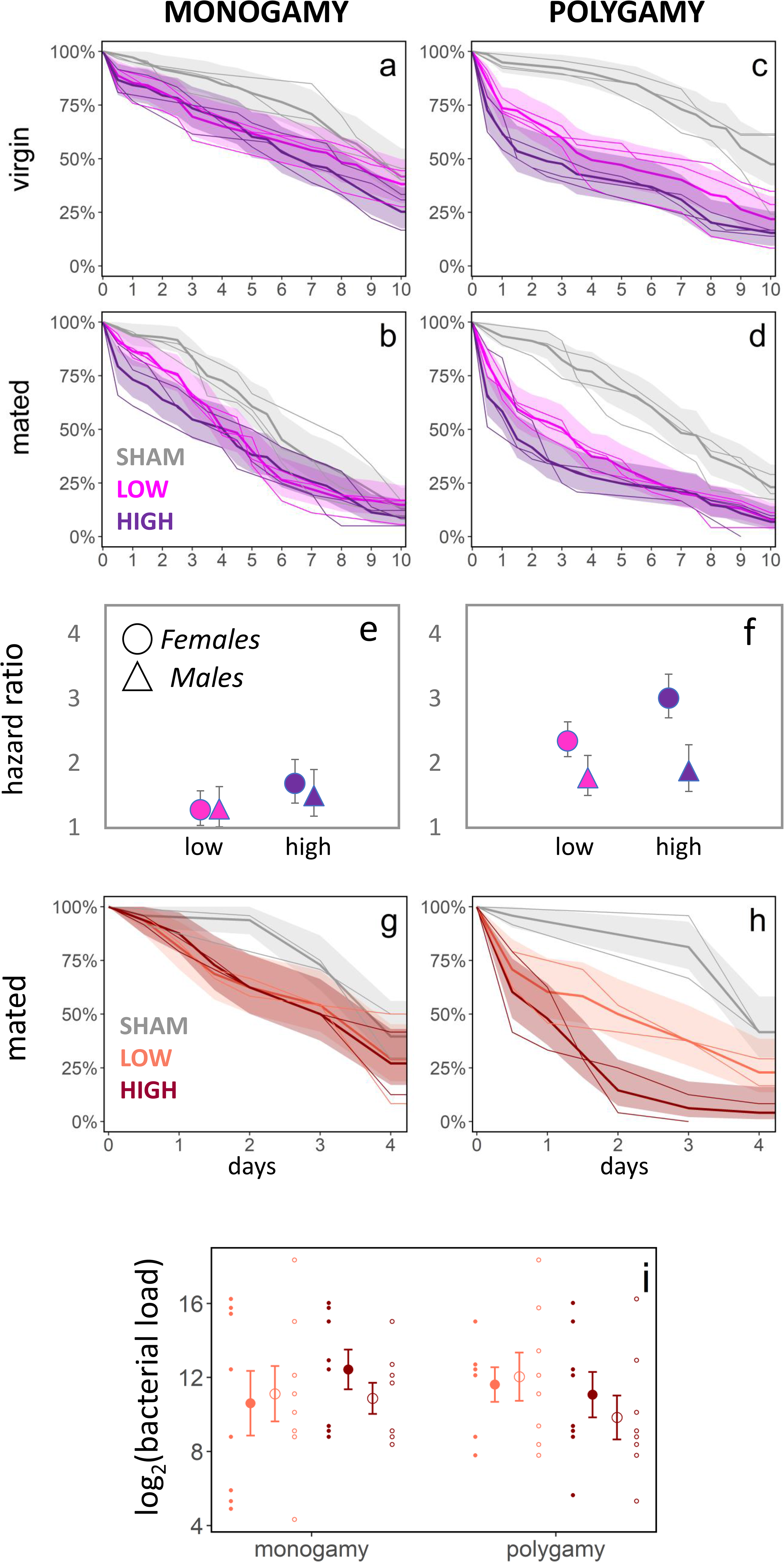
Microevolutionary change in tolerance to bacterial infection during experimental evolution under alternative mating regimes. Response to bacterial infection was estimated by the change in mortality rate between individuals infected with two doses of bacteria and a sham control. When infected with the gram-positive bacteria *B. thuringiensis*, monogamous females **(a, c)** had significantly higher survival under infection compared with polygamous females **(b, d)**, while virgin **(a, b)** and mated **(c, d)** females had similar responses. Shown are survival curves for each replicate evolution line (thin lines) together with mean survival (thick line) and 95% confidence limits (shaded area) based on all three replicate lines per regime and mating treatment. Virgin males (triangles) from monogamous **(e)** and polygamous **(f)** regimes did not show the strong differences seen in virgin females (circles), resulting in an apparent increase in sexual dimorphism in response to infection in the polygamy regime (compare panel **e** and **f**) (means ± 1SE; lower dose = 1.0 OD, higher dose = 2.0 OD for females and 2.5 OD for males). When mated females were infected with the gram-negative bacteria, *P. entomophila*, which allowed assaying of in vivo bacterial counts in infected individuals, monogamous lines **(g)** again showed higher survival under infection compared with polygamous lines **(h)** (lower dose = 0.5 OD, higher dose = 1.0 OD). **(i)** Counts of bacterial loads in females 12h post infection showed that difference in survival were likely not due to more efficient clearance of bacteria in monogamous lines. Means ± 1SE per replicate line (two lines used per regime and dose) and individual estimates per assay.

To gauge the generality of these results, and to further investigate whether the higher survival of monogamous females under bacterial infection was due to more efficient clearing of the bacterial infection (greater resistance), or because they were better at withstanding it (greater tolerance) (94), we infected once-mated polygamous and monogamous females with the gram-negative bacteria *Pseudomonas entomophila* using the same protocol as described above. The *P. entomophila* strain used is resistant to the antibiotic ampicillin. This allowed us to screen a subset of females collected 12h post start of infection exclusively for *P. entomophila* by culturing female cell tissue on Luria agar plates with ampicillin. Again, females from the polygamy regime showed higher susceptibility to bacterial infection (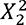 = 16.6, P < 0.001, total n = 288, Figure 4g,h, SI Tables 4d,e). However, there was no significant difference in bacterial load among the evolution regimes (P_MCMC_ > 0.2, n samples = 63, n females = 189, Figure 4i, SI Table 4f), suggesting no large differences in the ability of females to clear the bacterial infection. This last result does not support an allocation trade-off between the production of PO and AMPs, and may be more consistent with increased mortality due to toxic secondary metabolites resulting from overexpression of the proPO activating cascade by polygamous females (68,75–77) (Figure 1a). However, more work is needed to pin-point the exact mechanism underlying the differential mortality among monogamous and polygamous females. More generally, our results are consistent with the hypothesis that sexual conflict and harmful mating can lead to increased vulnerability to infection in females as a result of sex-specific trade-offs between different components of immunity (23,44,52).

### Correlated evolution between female PO activity and male genital morphology

We explored whether macroevolutionary transitions in sexual dimorphism in immunity could be driven by the evolution of mating interactions and the harmful morphology of male genitalia in this group of insects (23). We measured PO activity in virgin males and females of 12 species of seed beetles. There was pronounced sexual dimorphism and female-limited expression in many species (SI Figure 5a). To quantify harmfulness of the male genitalia in each species, we asked two expert and ten naïve biologists to rate pictures of male genitalia for the perceived harm they cause in the female reproductive tract (SI Figure 5b). Importantly, earlier work has shown that male harm assayed in this manner correlates positively with the amount of scarring that occurs in the female copulatory tract after mating (23). Species differences explained 61% of the total variation in rater scores and scores were highly correlated between experienced and naïve raters (r = 0.83), suggesting that raters generally agreed on the classification of male harm. Female and male PO activity, as well as male harm, showed moderate phylogenetic signals (Blomberg’s K = 0.68, 0.52 and 0.54, respectively (95)). Hence, we applied a phylogenetic generalized least squares regression (PGLS) based on species means using the ape package (96) for R, accounting for phylogenetic dependencies using Ohrstein-Uhlenbeck estimation and an extant seed beetle phylogeny (97, 98). There was significant positive covariance between male harm and female PO activity (α = 6.70, standardized slope = 0.83, df_12,10_, P < 0.001, SI Table 5a). Moreover, the covariance between male harm and male PO activity was not significant and opposite in sign (α = 2.92, standardized slope = −0.57, df_12,10_, P = 0.08, SI Table 5b). These analyses, together with our experimental findings, implicate sexual conflict as a driver of macro-evolutionary divergence in sexual dimorphism in immunity (Figure 5).

**Figure 5:**
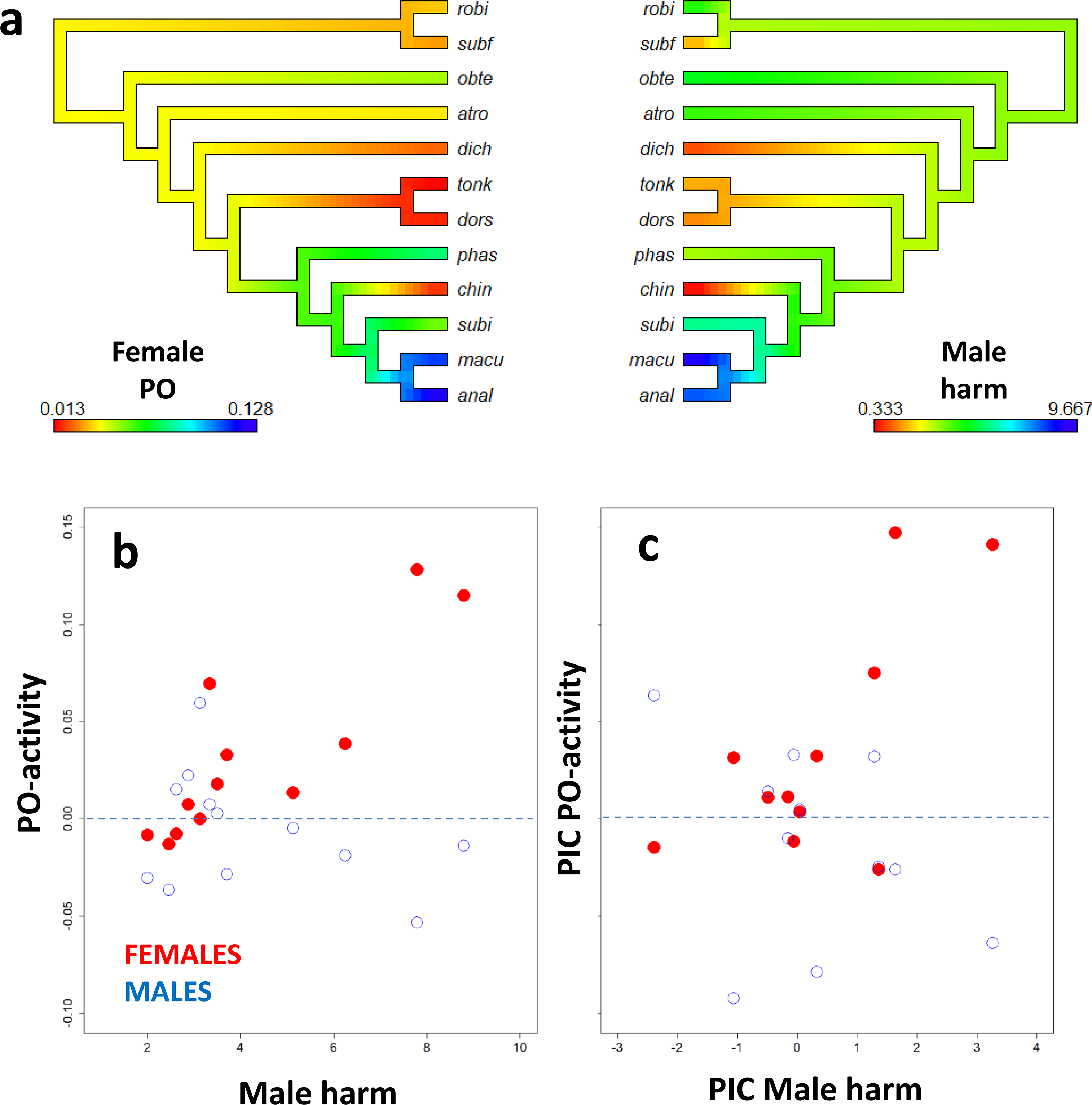
Phylogenetic covariance between harmfulness of male genital morphology and PO activity in virgin male and female seed beetles. **(a)** Female PO activity and the harmfulness of male genitalia mapped on the phylogeny of the 12 species used. Scores are given by color from blue (high harm/PO) to red (low harm/PO). Lower panels show correlations across species between male harmfulness and male (blue open) and female (red closed) PO activity, shown as **(b)** raw tip data and **(c)** phylogenetic independent contrasts (PICs). Standard errors around each species’ mean were typically of the magnitude ∼0.02 for male and female PO activity, and ∼0.6 for male genital morphology. The y-axes of **b)** and **c)** are scaled to have the same range. Species codes represent robi = *Amblycerus robinae*; subf = *Zabrotes subfasciatus*; obte = *Acanthoscelides obtectus*; atro = *Bruchidius atrolineatus*; dich = *Bruchidius dichrostachydis*; tonk = *Megabruchidius tonkineus*; dors = *Megabruchidius dorsalis*; phas = *Callosobruchus phaseoli*; chin = *Callosobruchus chinensis*; subi = *Callosobruchus subinnotatus*; macu = *Callosobruchus maculatus*; anal = *Callosobruchus analis*.

## Discussion

Sexual selection can result in increased male harm to females during mating (22,29,32), either through direct injury or infection with pathogens, and this should in theory favour increased female investment in immunity when female lifetime reproductive success is elevated by increased longevity (5,22–24,27,35,39). Here, we provide a suite of experimental and comparative data collectively showing that sex-differences in immunity can be modulated by sexual conflict in a species where costs of mating are conspicuous. This conclusion is based upon observations of (1) sex-biased expression of genes in the proPO activating cascade (Figure 1), (2) a female-bias in PO activity which is substantially higher than what is typical in insects, (3) female-limited phenotypic plasticity in PO activity in response to mating (Figure 2), (4) female-limited microevolutionary changes in immunity traits in response to experimental manipulation of the mating system and hence sexual conflict (Figures 3 & 4), and (5) correlated evolution between male genital morphology and female PO activity across species (Figure 5).

While previous studies have quantified female immune responses post mating (5,5,22,23,45,50,53,99,100), it often remains unclear whether male harm via genitalia or ejaculatory compounds (i.e. sexual antagonism) drive such responses, or whether they represent independent female optimization of the trade-off between current and future reproduction (5,22,23,27,49,101). Here, we directly manipulated the level of sexual selection and conflict, which is relatively well understood in *C. maculatus* (e.g. (47,48,59,62,102–105)), and found a clear female-limited PO response, while no correlation between female reproductive investment and PO activity was detected. Hence, our data point to male harm inflicted during mating as the driver of female PO investment. In this system, the inflicted harm by a male on his female mating partner is positively correlated to his success in sperm competition (29), presumably because seminal fluid substances (66) that benefit males in sperm competition (62) pass more rapidly into the female body if the copulatory duct is ruptured (32). However, these wounds may leave females at a risk of systemic infection with pathogens (36), suggesting a need for healing these injuries via a PO-mediated, potentially costly (68,73,76,77), reaction.

We hypothesized that these effects could have consequences for female susceptibility to infections unrelated to mating via trade-offs between PO activity and other components of immunity in the prophenolixidase activating cascade, such as the production of AMPs (Figure 1). This prediction was offered correlative support by the observation of increased susceptibility to bacterial infection in females from the polygamous mating regime (Figure 4). However, our results do not allow us to confidently distinguish between several, mutually inclusive, hypotheses regarding the exact mechanistic basis for the increased mortality of polygamous females. We did not find a difference in bacterial load between polygamous and monogamous females infected with the gram-negative bacteria *P. entomophila*, suggesting no large differences in the ability to clear infection and therefore also no large differences in the production of AMPs used to fight bacterial infection; a result that does not support a trade-off between the production of AMPs and PO in polygamous females. The result from this experiment is hardly conclusive, however. Another possibility is that the need for high PO activity in the reproductive tract of polygamous females led to a harmful “overactivation” of the proPO activating cascade upon bacterial infection and the simultaneous need for AMP production (e.g. (106)). Indeed, while such overactivation could mask allocation trade-offs by attending the dual need of producing PO and AMPs, it may have caused an inflammatory response with increased mortality of polygamous females as a result. The proPO activating cascade can have detrimental immunopathological consequences via the production of toxic secondary metabolites and needs to be strictly regulated (76), and severe bacterial infection can kill the organism also via side-effects of excessive melanization (68, 73). Future experiments are needed to pin-point the exact mechanistic basis underlying our results, preferably including detailed measures of tissue-specific immunity responses as we here measured whole-body samples.

Interestingly, polygamous females suffered increased mortality when infected with gram-positive and gram-negative bacteria, with only the former considered important in activating immune responses via TOLL, while the latter is thought to elicit immune responses mainly via the Imd pathway (78), which does not have a clear connection to the proPO activating cascade. This result might thus speak against immunity trade-offs between components in the proPO activation cascade as a general mechanism explaining the observed differential mortality between polygamous and monogamous females. However, several studies on invertebrates have now demonstrated cross-talk between the TOLL and Imd pathway ((55,78,107–109)), and that the proPO cascade can be readily activated by gram-negative bacteria (74,109,110). Indeed, TOLL has even been directly implicated in regulating sexual dimorphism in immunity to gram-negative bacteria in fruit flies (111), suggesting that the consistent difference in mortality of monogamous and polygamous females infected with the gram-positive *B. turingiensis* and gram-negative *P. entomophila* may yet be rooted in differential usage of the proPO activating cascade.

Male reproductive success in polyandrous mating systems is typically maximized by a shift towards current reproduction in the female mating partner, as this would increase the likelihood of the male siring a larger fraction of the offspring produced by the female (5,16,23,24,44). These ideas predict that males should evolve to manipulate females to invest in current reproduction at the expense of reduced immunity and longevity (22, 23). In line with these predictions, males with longer genital spines, that inflict more harm during mating, sire more offspring in *C. maculatus* (59, 62) and seem to stimulate female fecundity (*unpublished data*). Moreover, the male ejaculate regulates female immunity post mating in *Drosophila*, guppies, mice and humans (44,45,51,52,112,113), although it often remains unclear to what extent the effects are detrimental or beneficial to the female overall (5,22,23,27,114). It has even been suggested that males may gain fitness benefits by transferring sexually transmitted diseases that trigger shifts in female allocation towards current reproduction (21, 115), but this possibility lacks empirical support (116). In other insects, female PO either increases or decreases post mating and it has been suggested that in species where mating downregulates female PO activity, males corrupt the female immune function (23). While our results do not refute this hypothesis, they are also consistent with *C. maculatus* females being “primed” for harmful mating and that PO activity in females initially decreases post mating as a result of wound healing but is then quickly restored. Such female anticipatory immunity activation has been observed in *Drosophila* (117, 118) and bed bugs (119).

## Conclusions

When mating rate affects both sexual dimorphism in immunity and infection rates, this can result in intricate eco-evolutionary dynamics with demographic consequences for both host and pathogen (5,16,21–25,27). Our study suggests that sexual conflict over mating rate can drive sexual dimorphism in immunity and that allocation to different components of immunity may play an important role in mediating effects of mating on females. In *Drosophila*, mating increases immune responses in reproductive tissue, and in most insects mating decreases general immunity, but causality typically remains unclear (22, 23). Our results imply that baseline PO activity decreases in *C. maculatus* females as a genetic response to the alleviation of sexual conflict and harmful mating. Moreover, monogamous females, that evolved a reduced investment in PO activity relative to naturally polygamous females, showed an associated evolutionary increase in tolerance to bacterial infection in abdominal tissue adjacent to the reproductive tract, effects not seen in their conspecific males. This suggests that sex-specific trade-offs determine the mosaic of immune investment and that sexual selection and conflict affect the economics of these trade-offs. This complexity may explain some of the discrepancies found in the literature concerning female immune responses to mating (reviewed in: (5,23,27)) and motivates further explorations of the selection pressures affecting sexual dimorphism in immunity.

## Methods

### Study populations

*Callosobruchus maculatus* females lay eggs on seeds and larvae burrow into the seed where the entire development occurs. Beetles emerging from seeds are reproductively mature and require neither water nor food to reproduce successfully (e.g. (120, 121). Adults typically die 7-14 days after emergence in the absence of food or water (e.g.(122). All experiments used beetles originating from a genetic stock that was originally sampled in Lome, Togo, in 2010, and subsequently maintained as 41 isofemale lines in the laboratory to maintain the genetic variation present in the original population (123), before being mixed into a large, outbred, and genetically diverse experimental population (N ∼500). This genetic stock has been used in quantitative genetic designs (e.g. (102,123–125), artificial selection experiments (126), and experimental evolution (63,88,89) to demonstrate substantial sex-specific standing genetic variation in behavior, morphology, life-history and life time reproductive success, as expected given that the lines originate from the center of the species range (127).

The experimental evolution lines used to study the effect of the mating system on the evolution of sexual dimorphism in immunity are thoroughly described in (63, 88). In brief, the lines were maintained under standard temperature (29°C), humidity (50%RH) and light cycle (12L: 12D), and were reared on the preferred host plant (127) *Vigna unguiculata* (black-eyed bean). There are three replicate “Monogamy” lines, three “Polygamy” lines and two replicate “Male-limited” lines. Effective population size for the lines in each regime was kept approximately equal (N_e_ ≍ 150; N_Male-limited_ = 200, N_Monogamy_ = 246, N_Polygamy_ = 300) and the number of beans provided as egg laying substrate in each regime was standardized to give the same, relatively low, juvenile density (2-4 eggs/bean) to minimize (and equalize) larval competition (63). To implement the different regimes, selection was only applied for the first two days of adult life. However, the reproductive output over these first days typically corresponds to half of the total lifetime reproductive output (D. Berger, unpublished data). The regimes show differences consistent with generally positive effects of sexual selection on genetic quality in terms of increased female reproductive success and population productivity in polygamy lines relative to monogamy lines at generations 16 and 20, respectively (63). They also show differences in sexually selected male pre- and post-copulatory traits (88, 89).

### Expression of genes involved in the proPO activating cascade

To assay the effects of sex and mating status on the expression of relevant genes, we used data previously published in (128). Briefly, RNA sequencing (Illumina TruSeq) was used to test for sex differences in gene expression in virgin and mated age-matched beetles, separately for reproductive and non-reproductive tissues (i.e. abdomen and head & thorax, respectively). In the mating treatment, RNA was extracted 24h after mating. We pooled six individuals of each sex, tissue and treatment and replicated these pools three times. The transcriptome was assembled *de novo* (129), and differential expression analysed using edgeR, as described in(128). The candidate PO genes were detected using BLAST (tblastn search in the TSA database for *C. maculatus*, using the protein sequences as query) and here we report the ones with a significant sex difference in expression (with a false discovery rate adjusted p-value < 5%) in the virgin beetles in either tissue category.

### Phenoloxidase assays

Individual beetles were homogenized by 20 seconds of grinding with a pestle in an Eppendorf tube containing 20 µl Phosphate Buffered Saline (PBS). Samples were kept on ice until centrifuged at 17g for 10 min at 0°C, and the supernatants (10 µl) were stored at −80°C prior to the assay of PO activity. The frozen homogenates were analysed by an investigator uninformed of the samples’ identity and treatment affiliation, i.e as blind tests. Due to the small volume of each sample and high background due to the crude protein extract, the assay was first developed and optimized to ensure that proper enzyme kinetics were at hand, and phenylthiourea could completely block the activity (see Supplementary Information 6). In preliminary experiments the beetle homogenate was preincubated with curdlan (a β-1,3-glucan), trypsin or chymotrypsin to fully convert all zymogenic proPO to the active enzyme PO before assay of enzyme activity. However, the frozen homogenates did not show any increased PO activity after activation, indicating that the preparation method such as freezing at −80°C had converted all proPO into active enzyme PO (Supplementary Information 6). Dopamine, L-Dopa and 4-methylcathecol+hydroxyproline ethyl ester were each tested as substrate for *Callosobruchus* PO, and dopamine was shown to be the most efficient substrate and was used in the further experiments (rough estimates of Km in this crude homogenate for L-dopa Km≈ 6.3 mM, and for dopamine Km ≈ 0.2 mM, while 4-methylcathecol + hydroxyproline ethyl ester as substrate did not show linearity). For the experimental samples, six samples of beetle homogenate at a time were randomly chosen and thawed. After thawing, each individual beetle homogenate (3 µl) was incubated together with 7 µl PBS and 50 µl dopamine [10 mM in H_2_O] at 22°C. The reaction proceeded for 15 minutes after which 60 µl H_2_O was added to terminate the reaction and after centrifugation at 16000 x g for 1 min the absorbance at 420 nm was recorded. The enzyme assay was first developed to ascertain zero order kinetics, and due to the crude source of enzyme individual blank controls (without substrate) had to be measured before and after the reaction for each sample. This blank control was assayed containing 3 µl beetle homogenate, 7 µl PBS and 50 µl H_2_O, and was incubated and measured as the samples above. The enzyme activity is expressed as increase in absorbance at 420 nm per minute in the focal sample relative to its blank control (ΔA420/min).

### Sex-specific ontogenetic regulation of phenoloxidase activity

The eggs laid by the females in the mating status experiment (below) were followed through ontogeny. We sampled a total of 20 final instar larvae, 20 pupae and 14 adults. Larvae of *C. maculatus* could not be sexed. Pupae were sexed by abdominal morphology, for a total of 10 male and 10 female pupae. Virgin adults were collected as virgins within 0-36 hours post emergence. All individuals were weighed and measured for PO activity. We analysed differences between developmental stages by adding mass of the tissue analysed as a covariate in an ANCOVA. As we could not determine the sex of larvae, we performed one model that averaged effects across the sexes and one model where we excluded larvae and could retain sex. Both models showed significant differences between life stages.

### Female phenoloxidase activity in response to mating

We used males and females from the Lome base population, reared at standard conditions. All adults were virgin and between 24-48 hours old at the start of the experiment. On day one, 120 females were individually placed in small 30mm diameter petri dishes together with two males, in three separate bouts (40 females at a time). Matings were observed and mated females were immediately removed and placed into a 90mm diameter petri dish containing black eyed beans allowing females to oviposit. In total, 114 of the 120 females mated successfully over an observation period of 20 minutes per bout. A random set of 35 of these females were assigned to treatment 100 (mating on day one and then reproduction in isolation until being measured for PO activity on day three). The rest of the females were given the opportunity to mate on day two and day three, but all females did not mate on all days. This resulted in four treatment groups; 100, (mated on day 1 only), 110 (mated on day 1 & 2), 101 (mated on day 1 & 3) and 111 (mated on all days). Approximately two hours after the final mating on day three, all females were weighed and then measured for PO activity as described above. Measuring PO activity is time-consuming, and since preliminary analyses of the first batch of females suggested sufficient power to detect effects of mating status (see Figure 2), all females were not measured. The following sample sizes were attained for each treatment; 100: 15, 110: 7, 101: 13, and 111: 23 females. The treatment groups described above from the first experiment represent non-random samples of females, as not all females can be made to remate on a given day. We therefore conducted a second experiment with random samples of 100 (n = 15) and 001 (n = 15) females to confirm the main result from the first experiment. We counted the number of adult offspring produced by each female over the 48h of egg laying in the first experiment. We analysed the effect of mating status and number of offspring produced, including their interaction, on female PO activity in an ANCOVA. Female body mass at the time of homogenization was included as a covariate.

To determine whether female PO is allocated to eggs, 10 matured eggs per female were dissected out from 25 virgin females for a total of five samples containing 50 eggs each (corresponding to approximately 50% of the lifetime production of eggs of a single female). Samples were weighed and then subjected to the same crushing and centrifuging protocol as the mated females before being frozen at −80 °C and later measured for PO activity.

### Experimental evolution of phenoloxidase activity under alternative mating regimes

The experiment was performed following 27 generations of experimental evolution and one subsequent generation of common garden (polygamy) selection through standard culturing to remove any potential influence of parental environmental effects. PO activity was measured in the whole body of single male and female beetles from two replicate lines from each mating regime (6 lines in total). To manipulate the reproductive status of the beetles, newly emerged virgin adults (0-48h old) were either placed together in 90mm diameter petri-dishes in groups of five males and five females that were allowed to reproduce (“Reproducing” treatment), or in petri dishes with 5 males and 5 females but individually isolated in aerated Eppendorf tubes (“Virgin” treatment). All petri dishes contained black eyed beans, so that all beetles experienced the olfactory stimuli of the host beans, but only reproducing females could oviposit on the beans. After 46h, individuals were weighed before being put through the protocol to measure PO activity (see above). Beans from the mating treatment were stored until adult offspring emerged. Offspring were frozen and 20°C and later counted to estimate allocation to reproduction in all regimes. We set up the experiment in two separate batches one week apart in time, with each batch containing one replicate line of each evolution regime. We analysed differences among evolution regimes and mating treatments in Bayesian mixed effect models implementing Markov chain Monte Carlo simulations using the MCMCglmm package (90) in R (91). We ran separate models for males and females as PO activity was virtually undetectable in males. Evolution regime and mating treatment, including their interaction, were added as fixed effects and body mass was added as a covariate to control for the amount of tissue analysed as we used whole-body samples. We first tested for presence of a higher order interaction between mating status and evolution regime, which was non-significant and removed. We then evaluated significance of main effects by comparing the posterior distributions of marginal means for two groups in a given comparison (e.g. comparing mean PO activity of monogamous and polygamous females, averaged over the two mating statuses). In follow-up analyses we also assessed interactions between female body mass and mating status and evolution regime (to test for condition-dependence of PO activity; see Results). We blocked out effects of batch by adding it as a fixed effect. Similarly, we also blocked out the potential effect of freezing some individuals before homogenizing samples, something that had to be done for logistic reasons. Replicate line crossed with mating treatment, and adult mass when appropriate, were always included as random effects when estimating effects of evolution regime on PO activity. We used weak and unbiased priors for the random effects and ran models for 3,000,000 iterations, preceded by 100,000 burn-in iterations that were discarded, and stored every 3,000th iteration (thinning), resulting in 1,000 uncorrelated posterior estimates of the fixed effects upon which we calculated Bayesian P-values and 95% credible intervals. Prior specification and MCMC settings were the same for all models (exemplified in Supplementary Table 3c).

### Evolution of the response to bacterial infection

At generation 50, we collected beetles from each of the three replicate populations of the Monogamy and Polygamy regime and then maintained them under common garden conditions (natural polygamy) for one generation to minimize environmental parental effects. To measure evolved vulnerability to a bacterial pathogen, we first isolated 2-day-old experimental virgin females from each of the lines and paired them individually with a single male from their own line for 5 hours. Simultaneously, we also collected another subset of females that were held as virgin throughout the experiment. On day three post eclosion, we infected females with a strain (DSM 2046) of the entomopathogenic gram-positive bacteria *Bacillus thuringiensis,* described in (130). Beetles were first anesthetized with carbon-dioxide and then pricked at the lateral side of the lower abdomen, using a 0.1mm minutien pin (Fine Science Tools) dipped in overnight bacterial suspension of 1 OD or 2 OD (subcultured from an overnight culture of the bacteria). We performed sham infection with a pin dipped in sterile PBS solution. Following the start of the infection (or sham infection), we isolated females individually in 24 well-plates. We monitored individual survival at every 12 hours until 48 hours post infection and daily around 6pm for the next 8 days. Females still alive 10 days post infection (less than 30%) were right-censored in the subsequent survival-analysis. In a separate experiment, we also measured survival of infected 3-day old virgin males as described above.

At generation 54, we again collected mated females from two randomly selected replicate populations each of Polygamy and Monogamy and maintained them under common garden conditions. In the subsequent generation (Gen 55) we collected virgin females from each regime. We first mated two-day old females with a male from their own population. We then infected the females with a 0.5OD (52.5 ± 19.3 cells/beetle) or 1.0OD (237.5 ± 124.7 cells/beetle) solution of the gram-negative bacteria *Pseudomonas entomophila* using the same protocol as described above for *B. thuringiensis* (note that we could not calculate exact cell counts for *B thuringiensis* as the strain used lacked an antibiotic marker). Following the start of infection, we housed females individually in the 24 welled plates. Survival was first observed after 12 hours and a subset of beetles were taken out for bacterial load assay described below. We measured survival up to 120 hours after the start of infection.

The *P. entomophila* strain used is resistant to the antibiotic ampicillin. This allowed us to screen the females collected 12h post infection exclusively for *P. entomophila* by plating their whole-body extract homogenized in sterile PBS buffer on LB agar plates with ampicillin (0.1mg/ml), and subsequently counting bacterial cultures on the plates to estimate bacterial load. We first collected 3 surviving females 12hours after start of infection and transferred them to a micro-centrifuge tube. We then washed the three beetles together with 70% ethanol twice. Following the ethanol wash we again washed them with sterile water once. Subsequently, we added 90 µl of PBS and crushed the beetles together using a sterile micro-pestle. From this master-stock solution we made dilutions up to 10_-5_ in 96-welled plates. We spotted 3ul of each dilution on Luria agar plates with ampicillin. We kept the plates over night at 27°C and counted distinguishable *Pseudomonas entomophila* colonies. From the number of colonies, we calculated the bacterial load per female beetle and used that for further analyses. In total we calculated load for 8 samples per line and bacterial concentration. One sample was lost, resulting in a total of 63 samples (each based on 3 females). Analyses described in the Results and model specifications in Supplementary 4.

### Correlated evolution between PO activity and male genital morphology

We measured the PO activity of 5 virgin males and 5 virgin females of each of the 12 species (see Figure 5) using whole-body samples. All individuals were less than 48h old post adult emergence. As the species differ widely in body size, we modified the amount of PBS buffer added at homogenization to retain more equal concentration of tissue for all species in the original samples to be analysed for PO activity.

We used a modified version of the protocol of (23) to assess variation in the injuriousness of male genitalia. We first dissected out the male genitalium from 2 individuals per species. Each genitalium was photographed twice from complimentary angles to describe the 3D structure of the aedeagus (the intromittent apical part of male genitalia). This resulted in 48 photos of the 24 male samples. The two complimentary photos of each genitalium were placed together on a sheet and given a random ID to hide the species identity for raters. We asked 10 colleagues (evolutionary ecologists at our institution) to individually rate the 24 male genitalia on a scale from 0-10 in terms of the harm they predicted that the genitalia would cause inside the female reproductive tract during mating. Two of the authors of this study, with ample experience of sexual conflict theory and seed beetle biology (GA and JLR) also rated the genitalia (without knowledge of the recorded PO activity in the species, except for *C. maculatus*). The scores of naïve and experienced raters were highly aligned (see: Results), suggesting that the rating of male harmfulness was unbiased in terms of prior knowledge of the mating system. We extracted a mean score for predicted harmfulness for each of the 24 males based on scores from all 12 raters.

We analysed the covariance between harmfulness of the male genitalia and male and female PO activity based on species means across the phylogeny using phylogenetic least squares (PGLS) regression with Ohrstein-Uhlenbeck correction implemented in the ape package(96) for R (model specification and output in Supplementary Table 4). All variables were variance standardized in the analyses. Given the uncertainty of exact branch lengths, we set all branches to unit length. PO measurements were divided by the concentration of tissue in each sample prior to analysis.

## Declarations

### Author Contributions

IS performed all PO activity assays. EP performed experiments on mating status and ontogeny. JLR and EP collected data for species comparisons. JB and IMA maintained the selection lines. QC performed the experiments on PO activity in the lines. BB, DBa and IK planned and performed measures of responses to bacterial infection in the evolution lines. EI and AS performed the bioinformatic analyses. DBe analyzed all other data together with JB. QC, JB, EI and DBe produced the figures. DBe planned and conceived the study with considerable input from IS, GA and IK. DB wrote the first draft of the manuscript with input from all authors.

### Ethics Statement

This research was conducted according to national legislation. No permits are needed for research on invertebrates.

### Data accessibility

All data will be uploaded to the Dryad data repository upon final acceptance.

### Consent to publish

All authors and institutions have approved the submission

