## Supplementary Material for "Sexual conflict drives micro- and macroevolution of sexual dimorphism in immunity"

**Supplement 1:** Sex-specific regulation of gene expression related to the proPO activating system

**Supplement 2:** Mating status and phenotypic plasticity of PO activity in females

**Supplement 3:** Microevolution of PO activity.

**Supplement 4:** Responses to bacterial infection in the experimental evolution lines

**Supplement 5:** Macroevolutionary change and coevolution between male genital morphology and female PO activity.

**Supplement 6:** Optimization of PO-activity assays

#### Supplement 1: Sex-specific ontogenetic regulation of gene expression related to the proPO activating system.

**Supplementary Table 1:** Sex-biased gene expression in virgin and mated adults (see attached excel-file)

### Supplement 2: Mating status and phenotypic plasticity of PO-activity in females

**Supplementary Table 2:** ANOVA on the effects of mating treatment and egg laying on PO-activity.

```
lm(PO ~ weight+treatment*eggs,na.action=na.omit, data=mate)->mod2

Anova Table (Type II tests)

Response: PO
                  Sum Sq Df F value    Pr(>F)
weight         0.0005784  1  1.9198    0.1722
treatment      0.0168899  3 18.6867 3.229e-08 ***
eggs           0.0000252  1  0.0836    0.7737
treatment:eggs 0.0009753  3  1.0790    0.3667
Residuals      0.0147628 49
```

### Supplement 3: Experimental Evolution of PO-activity.

**Supplementary Table 3a:** Model specification and summary for analyses of effects of mating treatment and mating regime on adult weight.

```
prior1 = list(R = list(V = 0.001, nu = 0.002), G = list(G1 = list(V =
diag(0.001,4), nu = 5), G2 = list(V = 0.001, nu = 0.002)))

modAW <- MCMCglmm(log(aw) ~ treat*sex*Regime + date2,
random = ~us(treat:sex):line + dish, data = immunity,
family = "gaussian",prior = prior1, nitt=110000,slice=TRUE, burnin=10000,
thin=100, verbose = FALSE)

 DIC: -660.5602

 G-structure:  ~us(treat:sex):line

                        post.mean   l-95% CI u-95% CI eff.samp
M:sexf:M:sexf.line      2.716e-03  0.0002763 0.007739     1000
atV:sexf:M:sexf.line   -8.429e-05 -0.0029372 0.003751     1000
M:sexm:M:sexf.line     -2.319e-04 -0.0040262 0.002912     1000
atV:sexm:M:sexf.line   -8.202e-05 -0.0033890 0.003592     1000
M:sexf:atV:sexf.line   -8.429e-05 -0.0029372 0.003751     1000
atV:sexf:atV:sexf.line  2.486e-03  0.0003563 0.006842     1000
M:sexm:atV:sexf.line    1.590e-06 -0.0035273 0.003349     1000
atV:sexm:atV:sexf.line  3.451e-06 -0.0031724 0.003331     1000
M:sexf:M:sexm.line     -2.319e-04 -0.0040262 0.002912     1000
atV:sexf:M:sexm.line    1.590e-06 -0.0035273 0.003349     1000
M:sexm:M:sexm.line      2.963e-03  0.0003904 0.007973     1000
atV:sexm:M:sexm.line    4.087e-04 -0.0028881 0.004381     1000
M:sexf:atV:sexm.line   -8.202e-05 -0.0033890 0.003592     1000
atV:sexf:atV:sexm.line  3.451e-06 -0.0031724 0.003331     1000
M:sexm:atV:sexm.line    4.087e-04 -0.0028881 0.004381     1000
atV:sexm:atV:sexm.line  2.667e-03  0.0003275 0.007091     1000

               ~dish

     post.mean  l-95% CI u-95% CI eff.samp
dish  0.001222 3.774e-07 0.002554    571.2

 R-structure:  ~units

      post.mean l-95% CI u-95% CI eff.samp
units    0.0222  0.01979   0.0249     1000

 Location effects: log(aw) ~ treat * sex * evol + date2

                     post.mean  l-95% CI  u-95% CI eff.samp  pMCMC
Intercept(male-lim)  -5.600057 -5.687583 -5.516414   1146.7 <0.001 ***
treatVirgin           0.235430  0.130255  0.354929   1254.3  0.004 **
sex.m                -0.425792 -0.557175 -0.316267    739.9 <0.001 ***
evol.mono             0.086984 -0.033215  0.195206   1102.3  0.136
evol.poly            -0.004587 -0.115949  0.107136   1000.0  0.936
date2 batch2         -0.009607 -0.055843  0.030553   1000.0  0.636
treatV:sexm          -0.021025 -0.158023  0.128074   1000.0  0.802
treatV:evolmono       0.030475 -0.131892  0.208241   1070.6  0.698
treatV:evolpoly       0.061962 -0.095706  0.205128   1183.9  0.408
sexm:evolmono        -0.029061 -0.186639  0.145714    877.7  0.700
sexm:evolpoly         0.025689 -0.132726  0.192904   1000.0  0.730
treatV:sexm:evolmono -0.011103 -0.251075  0.207852    797.6  0.898
treatV:sexm:evolpoly -0.003060 -0.208250  0.204767   1000.0  0.930
```

**Supplementary Table 3b:** Analysis of differences in offspring production between evolution regimes in the 46h mating treatment using a mixed effect model based using REML estimation.

```
lmer(offspring/5 ~ evolution + date2 + (1|line), data=fec[1:37,]) -> mod


REML criterion at convergence: 214.8

Random effects:
 Groups   Name        Variance Std.Dev.
 line     (Intercept)  0.00    0.000
 Residual             33.81    5.814
Number of obs: 37, groups:  line, 6

Fixed effects:
              Estimate Std. Error t value
Intercept (male) 52.210      2.890  18.065
Evolution.mono  -3.813      2.383  -1.600
Evolution.poly   2.677      2.281   1.174
date2batch2      3.338      1.919   1.739

Analysis of Deviance Table (Type II Wald chisquare tests)

Response: offspring/5
           Chisq Df Pr(>Chisq)
evolution 7.4380  2    0.02426 *
date2     3.0242  1    0.08203 .
```

**Supplementary Table 3c:** Model specification and summary for analyses of effects of mating treatment and mating regime on male PO-activity.

```
prior1a = list(R = list(V = 1, nu = 10^-6),
G = list(G1 = list(V = 1, nu = 10^-6),G2 = list(V = 1, nu = 10^-6),
G3 = list(V = 1, nu = 10^-6),G4 = list(V = 1, nu = 10^-6)))

modPO1 <- MCMCglmm(POcorr*100 ~ treat*Regime*scale(aw) + date2 + frozen.alive,
random = ~line + treat:line + aw:line + treat:aw:line, data = imM,
family = "gaussian",prior = prior1a, nitt=3100000,slice=TRUE, burnin=100000,
thin=3000, verbose = FALSE)

 DIC: 799.2401

 G-structure:  ~line

     post.mean  l-95% CI u-95% CI eff.samp
line   0.05037 9.353e-08  0.09085     1000

               ~treat:line

           post.mean  l-95% CI u-95% CI eff.samp
treat:line   0.01622 1.069e-07  0.07501     1000

               ~aw:line

        post.mean l-95% CI u-95% CI eff.samp
aw:line    0.0164 2.07e-07   0.1064    1000

               ~treat:aw:line

              post.mean  l-95% CI u-95% CI eff.samp
treat:aw:line   0.04399 8.409e-08   0.2114   600.3

 R-structure:  ~units

      post.mean l-95% CI u-95% CI eff.samp
units    0.4959    0.324   0.6379    1000

 Location effects: POcorr * 100 ~ treat * Regime * scale(aw) + date2 +
frozen.alive

                                post.mean l-95% CI u-95% CI eff.samp pMCMC
Intercept (monogamy)             -0.92479 -1.34161 -0.49606   1000.0 0.002 **
treatmated                        0.29644 -0.19470  0.69702   1000.0 0.200
Regime.poly                       0.39662 -0.12329  0.96676   1000.0 0.118
Regime.male                      -0.03171 -0.56118  0.55625   1000.0 0.872
scale(aw)                         0.04250 -0.23234  0.33188   1000.0 0.768
date2batch2                      -0.04258 -0.39569  0.28392   1000.0 0.740
frozen.aliveN2                    0.33676 -0.13060  0.79397   1211.1 0.144
frozen.aliveY                     0.06616 -0.18781  0.33586   1000.0 0.628
treatmated:Regimepoly            -0.40597 -0.98814  0.26107   1000.0 0.182
treatmated:Regimemale             0.04196 -0.60943  0.65505   1000.0 0.892
treatmated:scale(aw)             -0.24605 -0.64610  0.16304   1000.0 0.272
Regimepoly:scale(aw)             -0.17127 -0.54120  0.22086   1000.0 0.368
Regimemale:scale(aw)             -0.17745 -0.53124  0.24447   1000.0 0.362
treatmated:Regimepoly:scale(aw)   0.19683 -0.38369  0.72463    832.3 0.526
treatmated:Regimemale:scale(aw)   0.31375 -0.24755  0.82759    701.9 0.268
```

#### Supplement 4: Responses to bacterial infection in the experimental evolution lines

**Supplementary Table 4a:** Cox-proportional hazards regression estimating the effect of infection with *B. thuringiensis* on survival of virgin and mated females from polygamous and monogamous regimes, following 50 generations of experimental evolution. Censused beyond 10 days post-infection.

```
cox_10 <- coxme(Surv(time = data2$hours, event = data2$censor) ~
regime*treatment*mating + block + (1|line), data = data2)


Cox mixed-effects model fit by maximum likelihood
  Data: data2
  events, n = 835, 1060
  Iterations= 5 27
                    NULL Integrated    Fitted
Log-likelihood -5330.719   -5248.14 -5248.031

                   Chisq    df p    AIC   BIC
Integrated loglik 165.16 14.00 0 137.16 70.97
 Penalized loglik 165.38 13.04 0 139.29 77.63

Fixed coefficients
                                         coef exp(coef)   se(coef)     z       p
regimeMO                          0.148866769 1.1605184 0.21984854  0.68 5.0e-01
treatment1 od                     0.879119736 2.4087784 0.19788298  4.44 8.9e-06
treatment2 od                     1.182189666 3.2615080 0.19656507  6.01 1.8e-09
mating                            0.635735994 1.8884115 0.19699346  3.23 1.3e-03
block2                            0.002521189 1.0025244 0.09242857  0.03 9.8e-01
block3                            0.027353935 1.0277315 0.08110524  0.34 7.4e-01
regimeMO:treatment1 od           -0.636142136 0.5293306 0.28486042 -2.23 2.6e-02
regimeMO:treatment2 od           -0.666092543 0.5137120 0.28018858 -2.38 1.7e-02
regimeMO:mating                   0.092003510 1.0963687 0.27608750  0.33 7.4e-01
treatment1 od:mating             -0.152276477 0.8587508 0.25400174 -0.60 5.5e-01
treatment2 od:mating             -0.246807326 0.7812912 0.25190300 -0.98 3.3e-01
regimeMO:treatment1 od:mating     0.130854710 1.1398022 0.36428569  0.36 7.2e-01
regimeMO:treatment2 od:mating     0.110071650 1.1163581 0.35843071  0.31 7.6e-01

Random effects
 Group Variable  Std Dev      Variance
 line  Intercept 0.0127639889 0.0001629194


Analysis of Deviance Table (Type II tests)
                        Df   Chisq Pr(>Chisq)
regime                   1  9.8152   0.001731 **
treatment                2 70.8506  4.121e-16 ***
mating                   1 63.6131  1.514e-15 ***
block                    2  0.1350   0.934710
regime:treatment         2 13.7176   0.001050 **
regime:mating            1  1.6283   0.201945
treatment:mating         2  1.1752   0.555654
regime:treatment:mating  2  0.1435   0.930786
```

**Supplementary Table 4b:** Bayesian mixed effects model estimating the effect of infection with *B. thuringiensis* on survival of virgin and mated females from polygamous and monogamous mating regimes, following 50 generations of experimental evolution. Binomial response estimated after 5 days of census, at which point 51.8% (549/1060) of all females, including controls, were still alive. Interactions with P>0.2 were removed from the model presented below.

#model specification

```
prior_line = list(R = list(V = 1, fix = 1),
G = list(G1 =  list(V = 1, nu =10^-6), G2 =  list(V = 1, nu =10^-6)))

MCMC_120 <- MCMCglmm(censor120 ~ mating + regime*treatment,
+  random = ~  line + treatment:line,
+  rcov = ~ units, data = data2,
+  family = "categorical", prior = prior_line, nitt=550000,
+  slice=TRUE, burnin=50000, thin=500)
```

#model output

```
 Iterations = 50001:549501
 Thinning interval  = 500
 Sample size  = 1000

 DIC: 1326.432

 G-structure:  ~line

     post.mean  l-95% CI u-95% CI eff.samp
line    0.01392 9.763e-08   0.06755     1000

               ~treatment:line

               post.mean l-95% CI u-95% CI eff.samp
treatment:line    0.0054 1.43e-07  0.03166     1000

 R-structure:  ~units

      post.mean l-95% CI u-95% CI eff.samp
units         1        1        1        0

 Location effects: censor120 ~ mating + regime * treatment

                       post.mean l-95% CI u-95% CI eff.samp  pMCMC
(Intercept)               1.9563   1.4238   2.5020   1000.0 <0.001 ***
regimeMO                 -0.2939  -0.9984   0.4853   1000.0  0.452
mating                   -0.9125  -1.3224  -0.3956   1000.0 <0.001 ***
treatment1 od            -2.0138  -2.5720  -1.5103    843.6 <0.001 ***
treatment2 od            -2.4411  -3.0202  -1.8552   1000.0 <0.001 ***
regimeMO:treatment1 od    0.9421   0.1318   1.7124   1000.0  0.018 *
regimeMO:treatment2 od    1.2943   0.5221   2.0855   1000.0 <0.001 ***
```

**Supplementary Table 4c:** Cox-proportional hazards regression estimating the effect of bacterial infection on survival of virgin males from polygamous and monogamous regimes following 50 generations of experimental evolution. Censused over 5 days following infection, when 35% (176/270) of all males were still alive. There were significant effects of the bacterial infection (P = 0.021) and monogamous males died faster than polygamous males (P = 0.010), but there was no difference in the effect of infection between regimes (P = 0.63).

```
Cox mixed-effects model fit by maximum likelihood
  Data: data
  events, n = 176, 270
  Iterations= 9 48
                    NULL Integrated    Fitted
Log-likelihood -908.3705  -896.3209 -894.5783

                  Chisq   df          p   AIC    BIC
Integrated loglik 24.10 8.00 0.00220560  8.10 -17.26
 Penalized loglik 27.58 7.62 0.00042412 12.34 -11.83

Model:  Surv(time = data$hours, event = data$censor) ~ regime * treatment +
block + (1 | line)
Fixed coefficients
                                 coef exp(coef)  se(coef)     z     p
regimeMO                    0.6977231 2.0091728 0.3042473  2.29 0.022
treatment1OD                0.6146787 1.8490625 0.2953745  2.08 0.037
treatment2.5OD              0.6576819 1.9303126 0.2982778  2.20 0.027
block2                      0.2042124 1.2265587 0.2282020  0.89 0.370
block3                      0.3967714 1.4870159 0.2282114  1.74 0.082
regimeMO:treatment1OD      -0.3668521 0.6929121 0.3834369 -0.96 0.340
regimeMO:treatment2.5OD -0.2485621 0.7799214 0.3848408 -0.65 0.520



Analysis of Deviance Table (Type II tests)

Response: Surv(time = data$hours, event = data$censor)
                 Df  Chisq Pr(>Chisq)
regime            1 6.6288    0.01003 *
treatment         2 7.7733    0.02051 *
block             2 3.0237    0.22051
regime:treatment  2 0.9354    0.62644
```

**Supplementary Table 4d:** Cox-proportional hazards regression estimating the effect of infection with *P. entomophila* on survival of mated females from polygamous and monogamous mating regimes following 55 generations of experimental evolution. Censused beyond 5 days post-infection.

```
Cox mixed-effects model fit by maximum likelihood
  Data: data
  events, n = 209, 288
  Iterations= 14 87
                    NULL Integrated    Fitted
Log-likelihood -1077.392  -1041.207 -1039.683

                  Chisq   df          p   AIC   BIC
Integrated loglik 72.37 7.00 4.8972e-13 58.37 34.97
 Penalized loglik 75.42 6.43 5.6510e-14 62.55 41.04

Model:  Surv(time = data$hours, event = data$censor) ~ regime * treatment +
block + (1 | line)
Fixed coefficients
                                      coef exp(coef)  se(coef)     z      p
regimePolygamy                 -0.07289093 0.9297022 0.2928655 -0.25 0.8000
treatment0.5OD                  0.42975950 1.5368879 0.2530109  1.70 0.0890
treatment1OD                    0.47976262 1.6156908 0.2514013  1.91 0.0560
block2                          0.26041180 1.2974643 0.1873123  1.39 0.1600
regimePolygamy:treatment0.5OD   0.41907793 1.5205588 0.3562258  1.18 0.2400
regimePolygamy:treatment1OD     1.36338175 3.9093916 0.3511195  3.88 0.0001

Random effects
 Group Variable  Std Dev    Variance
 line  Intercept 0.12379893 0.01532618

Analysis of Deviance Table (Type III tests)

Response: Surv(time = data$hours, event = data$censor)
                 Df   Chisq Pr(>Chisq)
regime            1  0.0619  0.8034469
treatment         2  4.2609  0.1187816
block             1  1.9328  0.1644516
regime:treatment  2 16.5703  0.0002522 ***
```

**Supplementary Table 4e:** Bayesian mixed effects model estimating the effect of infection with *P. enthomophila* on survival of mated females from polygamous and monogamous mating regimes following 55 generations of experimental evolution. Binomial response estimated after 72h of census, at which point ca. 50% of all females, including controls, were still alive.

```
prior_line = list(R = list(V = 1, fix = 1),
+                   G = list(G1 =  list(V = 1, nu =10^-6), G2 =  list(V =
1, nu =10^-6)))

MCMC_3 <- MCMCglmm(censor72 ~ regime*treatment + block,
+                    random = ~  line + treatment:line,
+                    rcov = ~ units, data = data.frame(data),
+                     family = "categorical", prior = prior_line,
+                     nitt=1050000,
+                    slice=TRUE, burnin=50000, thin=500, verbose = FALSE,
+                    pr=F)


 Iterations = 50001:1049501
 Thinning interval  = 500
 Sample size  = 2000

 DIC: 328.3504

 G-structure:  ~line

    post.mean  l-95% CI u-95% CI eff.samp
line      1139 7.705e-08    10.32     2000

              ~treatment:line

              post.mean  l-95% CI u-95% CI eff.samp
treatment:line    0.2049 1.306e-07    0.977     1818

 R-structure:  ~units

     post.mean l-95% CI u-95% CI eff.samp
units         1        1        1        0

 Location effects: censor72 ~ regime * treatment + block

                              post.mean l-95% CI u-95% CI eff.samp pMCMC
(Intercept)                      1.0193  -2.0032   3.1227     1512 0.171
regimePolygamy                  -0.2294  -1.9480   3.6129     2000 0.425
treatment0.5OD                  -0.9935  -2.2559   0.2498     2000 0.113
treatment1OD                    -1.2189  -2.4438   0.1292     2000 0.072 .
block2                           0.6998  -1.7666   3.3902     2000 0.282
regimePolygamy:treatment0.5OD   -1.5373  -3.3340   0.3553     2000 0.098 .
regimePolygamy:treatment1OD     -4.1079  -6.3302  -1.8884     2000 0.001***
```

**Supplementary Table 4f:** Bayesian mixed effects model estimating the log counts of *P. enthomophila* in mated females from polygamous and monogamous regimes 12h post-infection.

```
prior_load = list(R = list(V = 1, fix = 1),

                  G = list(G1 =  list(V = 1, nu =10^-6), G2 =  list(V = 1,
                  nu =10^-6)))

MCMC_load <- MCMCglmm(logload ~ regime*treatment + block,
+                     random = ~  line + treatment:line,
+                     rcov = ~ units, data = data.frame(load),
+                     family = "gaussian", prior = prior_load, nitt=1050000,
+                     slice=TRUE, burnin=50000, thin=500, verbose = FALSE,
+                     pr=F)


 Iterations = 50001:1049501
 Thinning interval  = 500
 Sample size  = 2000

 DIC: 190.6991

 G-structure:  ~line

     post.mean  l-95% CI u-95% CI eff.samp
line     105.6 9.443e-08   0.7772     2000

               ~treatment:line

               post.mean l-95% CI u-95% CI eff.samp
treatment:line     0.133 1.23e-07   0.1376     2000

 R-structure:  ~units

      post.mean l-95% CI u-95% CI eff.samp
units         1        1        1        0

 Location effects: logload ~ regime * treatment + block

                           post.mean l-95% CI u-95% CI eff.samp pMCMC
(Intercept)                   3.1410   2.4566   4.2033     2000 0.012 *
regimePolygamy                0.3206  -0.7685   1.4896     2000 0.476
treatment1OD                  0.2361  -0.6107   0.9527     2000 0.509
block2                        0.2469  -0.8640   0.8081     2000 0.633
regimePolygamy:treatment1OD  -0.6714  -1.7594   0.5098     2000 0.204
```

### **Supplement 5: Macroevolutionary change and sexually antagonistic coevolution between male genital morphology and female PO-activity**

**Supplementary Figure 5a:** Macroevolutionary change in sexual dimorphism in PO-activity across seed beetle linages.

Species codes are: robi = *Amblycerus robinae*; subf = *Zabrotes subfasciatus*; obte = *Acanthoscelides obtectus*; atro = *Bruchidius atrolineatus*; dich = *Bruchidius dichrostachydis*; tonk = *Megabruchidius tonkineus*; dors = *Megabruchidius dorsalis*; phas = *Callosobruchus phaseoli*; chin = *Callosobruchus chinensis*; subi = *Callosobruchus subinnotatus*; macu = *Callosobruchus maculatus*; anal = *Callosobruchus analis*.

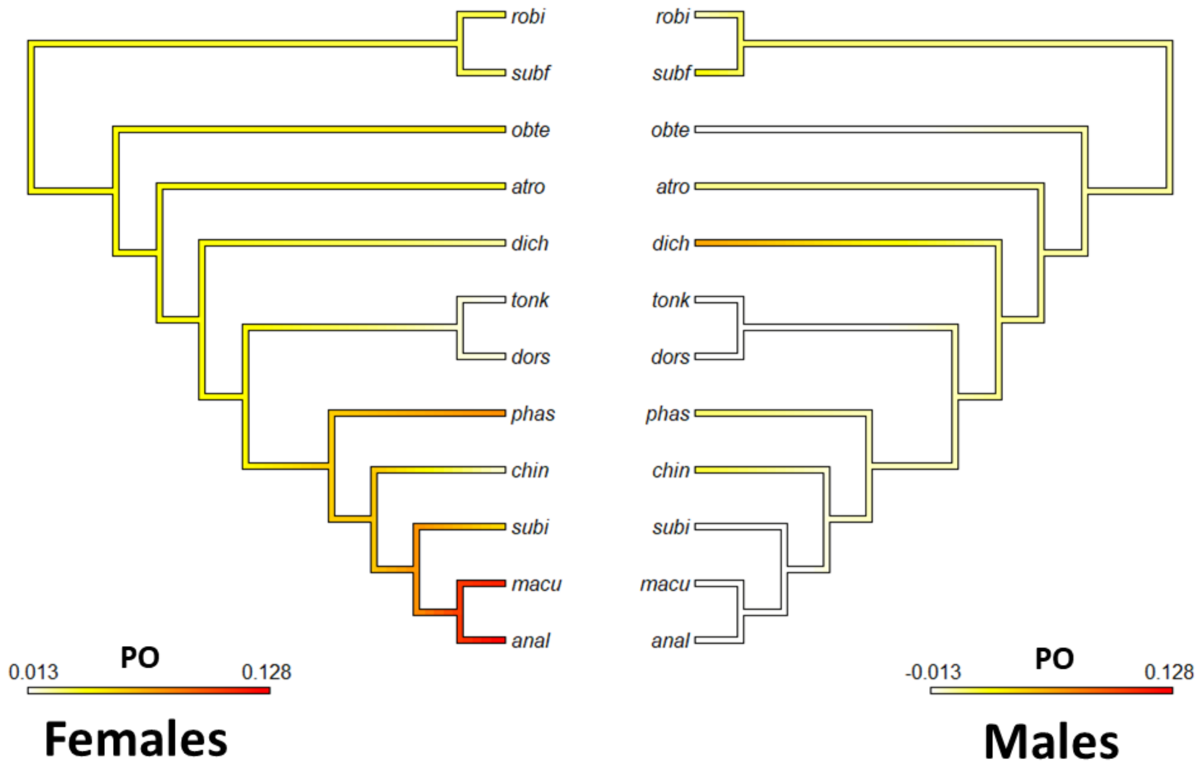

**Supplementary Figure 5b:** Photos of genitalia rated as least and most harmful.

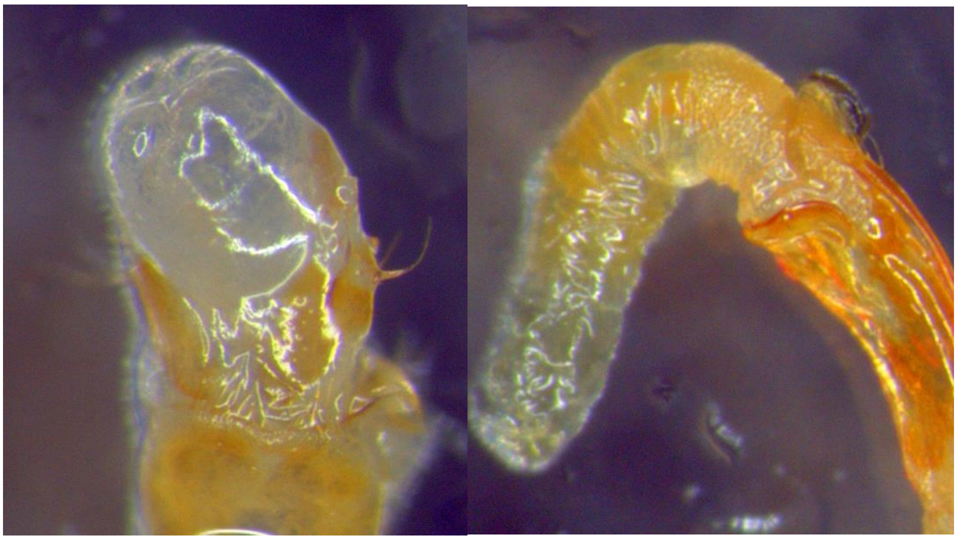

**Least harmful** (left: *Callosobruchus chinensis*; right: *Megabruchidius dorsalis*).

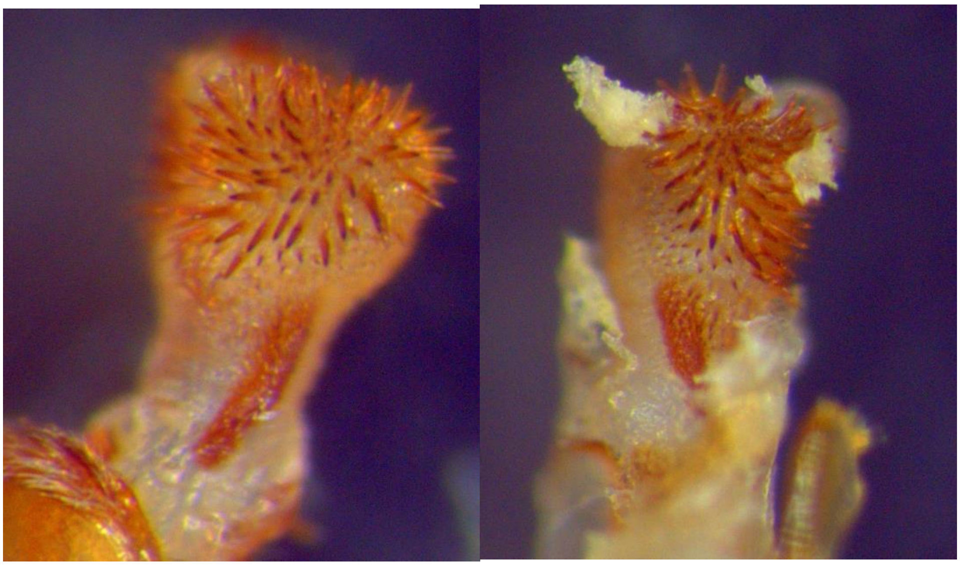

**Most harmful** (left: *Callosobruchus maculatus*; right: *Callosobruchus analis*).

**Supplementary Table 5a:** PGLS with Ornstein-Uhlenbeck correction between male genital morphology and female PO-activity.

```
Generalized least squares fit by REML
  Model: scale(PO.corr) ~ scale(genitalia)

       AIC      BIC    logLik
  27.04717 28.25751 -9.523586

Correlation Structure: corMartins
 Formula: ~1
 Parameter estimate(s):
   alpha
6.703784

Coefficients:
                      Value Std.Error   t-value p-value
(Intercept)      -0.0004046 0.1586664 -0.002550  0.9980
scale(genitalia)  0.8268925 0.1643090  5.032545  0.0005

Residual standard error: 0.5061646
Degrees of freedom: 12 total; 10 residual
```

**Supplementary Table 5b:** PGLS with Ornstein-Uhlenbeck correction between male genital morphology and male PO-activity.

```
Generalized least squares fit by REML
  Model: scale(PO.corrM) ~ scale(genitalia)
       AIC      BIC    logLik
  37.50322 38.71356 -14.75161

Correlation Structure: corMartins
 Formula: ~1
 Parameter estimate(s):
   alpha
2.918102

Coefficients:
                      Value Std.Error   t-value p-value
(Intercept)       0.0753423 0.3381249  0.222824  0.8282
scale(genitalia) -0.5679619 0.2901395 -1.957548  0.0788

Residual standard error: 0.9206185
Degrees of freedom: 12 total; 10 residual
```

### Supplement 6: Optimization of PO-activity assays

In order to test kinetics of the phenoloxidase assay, we prepared samples by mixing homogenates from three beetles, incubating as described in the Material and Methods, and followed absorbance at 420 nm each minute for 25 min to check linearity of the reaction. This experiment was repeated with three different homogenates, and the result is shown in the figure below. In addition, one pooled sample was preincubated with phenylthiourea (PTU) at 10 mM final concentration.

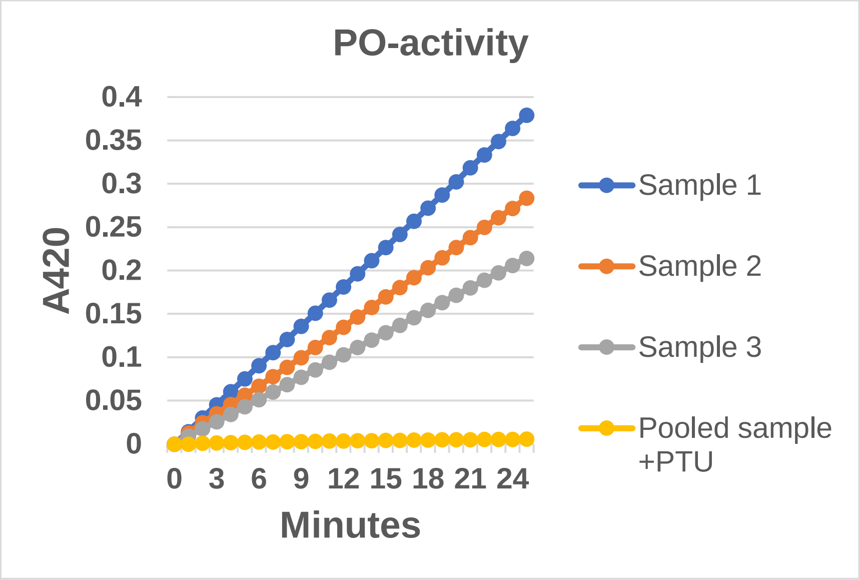


To test if all proPO was converted to PO in the frozen homogenates used in our experiments, the activity of pooled samples from three females were tested for activity after preincubation with curdlan (a β-1,3-glucan) at 1 mg/mL, or trypsin (0,1 mg/mL) or chymotrypsin (0,1 mg/mL). As shown in the table below, no additional activation could be observed after freezing the homogenates. We note that when assaying PO activity in fresh (unfrozen) samples, it is important to always test the presence of inactive proPO, since in a few fresh homogenates some minor increased PO-activity after preincubation with trypsin or chymotrypsin was found, indicating presence of intact proPO.

**Supplementary Table 6:** PO-activity after preincubation of frozen homogenate with different activators.

| | Sample A | Sample B | Sample C |
|---|---|---|---|
| Treatment | ΔA420/min | ΔA420/min | ΔA420/min |
| No activation | 0,0382 | 0,0429 | 0,0223 |
| Trypsin | 0,0368 | 0,0428 | 0,0217 |
| Chymotrypsin | 0,0375 | 0,0435 | 0,0197 |
| Curdlan | 0,0377 | 0,0433 | 0,0203 |
